# A farewell to EQ: A new brain size measure for comparative primate cognition

**DOI:** 10.1101/2021.02.15.431238

**Authors:** Carel P. van Schaik, Zegni Triki, Redouan Bshary, Sandra Andrea Heldstab

## Abstract

Both absolute and relative brain size vary greatly among and within the major vertebrate lineages. Scientists have long debated how larger brains in primates and hominins translate into greater cognitive performance, and in particular how to control for the relationship between the non-cognitive functions of the brain and body size. One solution to this problem is to establish the slope of cognitive equivalence, that is the line connecting organisms with an identical bauplan but different body sizes. Here, we suggest that intraspecific slopes provide the best available estimate of this measure. This approach was abandoned because slopes were too low by an unknown margin due to estimation error. We control for the error problem by focusing on highly dimorphic primate species with large sample sizes and fitting a line through the mean values for adult females and males. We obtain the best estimate for the slope of ca 0.27, a value much lower than those constructed using all mammal species, and close to the value expected based on the genetic correlation between brain size and body size. We also find that the estimate of cognitive brain size based on cognitive equivalence fits empirical cognitive studies better than the encephalization quotient (EQ), which should therefore be avoided in future studies on primates, and presumably mammals and birds in general. The use of residuals from the line of cognitive equivalence may change conclusions concerning the cognitive abilities of extant and extinct primate species, including hominins.

## Introduction

Although recent ecological approaches to comparative cognition have focused on linking performance in specific cognitive tasks to specific brain regions [e.g. Healy & Krebs 1996], traditionally comparative cognition has relied on a presumed link between some summary measure of cognitive performance and total brain size [Jerison 1973]. Scholars have therefore long been searching for a neuroanatomical measure of overall cognitive ability, both to compare living species and to estimate the cognitive abilities of extinct ones relative to their extant relatives. The most intuitive measure is a species’ brain size (or, especially for extinct species, the highly similar cranial capacity: Isler et al. 2008]. However, because brains also control numerous non-cognitive somatic functions, most researchers have agreed it cannot be used without controlling for these somatic functions.

Building on a century-long tradition, Jerison [1973] distinguished between the somatic and the cognitive brain functions and proposed we can estimate the portion of the brain dedicated to somatic functions, and so by subtraction arrive at the size of the cognitive portion. Jerison fitted regression equations to the brain size-body size data of all available species in many different mammalian lineages. Because this produced a regression a slope close to 0.67 in a log-log plot, he interpreted this as reflecting a fundamental physiological regularity linking brain size to the body’s surface area and thus the amount of proprioceptive inputs. This slope could therefore serve as the expected value of the somatic portion of the brain, and the deviation from the regression line as the estimate of the cognitive portion, i.e. cognitive brain size. Jerison then proposed the encephalization quotient (EQ), the ratio of a species’ actual brain size to its predicted brain size based on the clade-specific brain-body regression line, to capture its relative cognitive performance. The EQ has become a commonly used estimate of the cognitive abilities or intelligence of animal species. Other researchers have argued the slope is actually somewhat higher, linking it to metabolic turnover instead, and thus suggesting a slope of 0.75 [Martin 1981; Armstrong 1983], but did not question the fundamental EQ approach.

Most research on basic relationships between brain size, body size and cognitive performance has been conducted on mammals, and in particular primates. This research has produced various lines of evidence argue against the EQ approach. First, its use assumes that there is no correlation between body size and cognitive performance, but in practice there is a negative correlation between EQ and body size within mammalian orders (see below). We should therefore see that bigger mammal species show poorer cognitive performance than smaller relatives in the same order. However, the opposite appears to be the case [Rensch 1973], suggesting a lower value of the slope. Second, a lower slope would also be expected given the combined effect of two persistent macro-evolutionary trends: (i) brains have increased relative to body size, a phenomenon we can call the Lartet-Marsh rule [Jerison 1973], and (ii) body size did increase as well, a phenomenon known as Cope’s rule [Alroy 1998]. As a result, more recently evolved lineages will tend to have both larger body sizes and a larger brain sizes [e.g. Halley & Deacon 2017], which artificially inflates the slope of the regression through the total sample. Third, different mammalian lineages unexpectedly show different regression slopes of brain size on body size [Martin & Harvey 1985].

These inconsistencies reveal the fundamental flaw in the EQ approach [Deacon 1990; Striedter 2005]: it assumes that one process (be it proprioception or metabolic turnover) predominates in determining brain size to the extent that cognitive functions produce only minor deviations, so we can estimate their strength by taking ratios or residuals. Instead, we may expect a variety of processes to affect brain size, some of them lineage-specific, and each potentially varying in strength across lineages. Nonetheless, EQs continue to be used to compare mammalian taxa [e.g. Boddy et al. 2012], especially extinct ones [e.g. Grabowski et al. 2016; Benoit et al. 2019].

Although EQ approaches are inadequate, no universally accepted alternative has emerged. A major reason is that the notion of overall cognitive ability or performance remained vague, making it harder to produce an alternative measure. In fact, modern behavioral ecology generally assumes animals have bundles of domain-specific cognitive adaptations [Shettleworth 2010]. Consequently, “overall” cognitive performance does not exist and overall brain size does not necessarily provide any useful information when it comes to understanding the animal’s ecological niche or social organization. Instead, researchers relate the relative size of specific brain regions to specific cognitive challenges to look for patterns at that level in large comparative surveys. Examples include seed caching and recovery [Healy & Krebs 1996: Garamszegi & Eens 2004] or the incidence of feeding innovations [Timmermans et al. 2000] in birds.

Recently, however, brain size has regained standing as a relevant variable. First, along with the unquestioned selective increase or decrease of certain brain parts in response to specific pressures [Barton & Harvey 2000], mammalian brains are also organized in fundamentally similar ways across animals of varying sizes, even across lineages, with predictable allometric relationships among the sizes of the various brain regions [Finlay & Darlington 1995; Finlay et al. 2001]. Indeed, many cognitive activities correspond to concerted activity waves throughout the brain rather than being tied to one particular region [Park & Friston 2013], suggesting that a decrease or increase in one particular region under selection will affect the sizes of other parts and thus also the size of the whole brain [Barton 2006]. Second, an important consequence of cognitive abilities that tie together many brain regions is domain-general intelligence [Burkart et al. 2017]. This concept has historically been applied mainly to humans, and many, implicitly or explicitly, still consider it to be uniquely human. There is now mounting evidence, however, that at least among various mammals [Burkart et al. 2017] and birds [Ashton et al. 2018], domain-general intelligence can be recognized within species, and that different species vary considerably in the strength of this domain-general ability [Deaner et al. 2006; Reader et al. 2011]. The latter finding allows empirical tests of the predictive value of EQs or conceptually similar residuals. Broad comparative analyses of estimated domain-general cognitive abilities have shown that EQ is a poor predictor of these abilities [Deaner et al. 2007; Reader et al. 2011], and more limited analyses reached the same conclusion [Alba 2010; Rumbaugh et al. 1996]. Moreover, as noted above, most large-bodied species have greater cognitive abilities than expected based on their EQ values [Rensch 1973], which tend to decrease with body size within orders. In conclusion, these more direct, cognition-based tests of EQ-based approaches confirm that they do not predict cognitive abilities.

### Non-EQ approaches

These conceptual and empirical problems with EQ measures indicate that we need to control for body size in a different way. Some scholars have suggested that no control is needed at all [e.g. Rensch 1973; Striedter 2005; Byrne 1995], but most accept that the regulation of somatic functions requires at least some brain resources that are not also available for cognitive functions. Two main alternatives have been proposed. The first set of techniques is similar to the EQ approach, as it relies on broad interspecific comparisons. Portmann [1946, 1947] proposed that Galliformes are the most primitive birds and deviations from their interspecific regression equation should be used to estimate encephalization. Stephan [1960] proposed the same procedure for mammals, using the “basal insectivores” as the baseline. This produces a “progression index” for each species. However, due to various conceptual and statistical problems [Deacon 1990] it has found little application. This index also predicts primate cognitive performance only marginally better than EQ and far worse than brain size per se [Gibson et al. 2001; Deaner et al. 2007], although it is unknown how well it does for other mammals or birds.

Another interspecific approach uses ratios of different brain regions, assuming that one, the numerator, is responsible for the brain’s cognitive functions, and the other, the denominator, is responsible for its somatic functions [Krompecher & Lipák 1966; Passingham 1982]. This ratio then estimates the development of cognitive functions relative to expectation. The most popular measure is the Neocortex Ratio, the size of the neocortex relative to the rest of the brain [Dunbar 1992]. However, all ratio measures have the fundamental drawback that they lack a clear neurobiological justification [Deacon 1990; Deaner et al. 2000; Barton 2006], and the neocortex ratio was favored simply because it yielded the best correlation with the putative selective pressure [Dunbar 1992]. Like most other ratios, the neocortex ratio is clearly correlated with both overall brain and body size, as expected based on the fundamental brain allometries [Finlay & Darlington 2001; Halley & Deacon 2017]. Thus, its use depends entirely on the goodness of the fit with presumed selective pressures, risking circularity. If this fit varies across taxa, we cannot tell whether a poor fit reflects an imperfect neuroanatomical measure or different selective environments [Stout 2018]. Moreover, the neocortex ratio does not differentiate between monkeys and apes, which are known to differ in cognitive performance [Gibson et al. 2001]. Also, it does not always predict actual cognitive performance in the primate sample as strongly as other measures when we control for phylogenetic non-independence [e.g., Deaner et al. 2007]. We will therefore not further consider ratio measures (but see discussion).

### The slope of cognitive equivalence

The aim of this paper is to revisit the main remaining alternative: estimating the slope of cognitive equivalence. Its logic works as follows. We assume (1) that the cognitive performance of adult individuals within a species does not depend on their body size, and (2) that the intraspecific relationship between body size and brain size among adults estimates the extra amount of brain tissue required to sustain the additional somatic functions in larger individuals because there are no changes in bauplan and sensory-motor abilities. Therefore, assuming that intraspecific variation in adult size is based on selection on body size alone, the regression of brain size on body size should give us the slope of cognitive equivalence. If analyses conducted on a range of species converge on a similar slope, the cognitive brain size of a species can then be estimated as the residual of the actual brain size of that species from a general line with that slope and some informative intercept.

This is not a new idea. Many estimates of this slope from samples of conspecific individuals of known body and brain mass have been made in the past. However, they vary considerably. Summarizing previous work, Pilbeam & Gould [1974] noted values between 0.2 and 0.4. The range for individual mammal species with larger samples was somewhat tighter, with average values between 0.23 and 0.25 (e.g. Hemmer 1971; Röhrs 1986] for large intraspecific samples, but a median of 0.14 for primates [Martin & Harvey 1985] and 0.15 in a large recent analysis of birds and mammals [Tsuboi et al. 2018]. This considerable variability exists because empirical slopes within species are usually underestimated [Pagel & Harvey 1988] by varying margins. Thus, if we can remove this source of error, the this measure’s utility could be restored.

The variability in slope estimates can be explained as follows. The equation for the slope of a regression is [Lande 1985]:

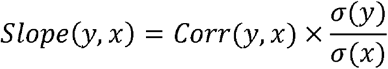

where corr(y,x) is the correlation between x and y, and *σ* is the standard deviation. Because virtually all mammals show determinate growth, there is little variation in body and brain size. As a result, variation due to measurement error can greatly affect the estimated slope of the brain-body relation within a set of adults. We usually estimate the independent variable, body size, by taking body mass, but there are good reasons to assume that this estimate varies considerably due to error, even within individuals. In contrast, the response variable, brain mass or endocranial volume (our estimate for brain size), varies far less. First, during periods of starvation, brains continue to receive the same energy flow as during times of plenty, whereas the rest of the body must make do with less. This phenomenon is known as brain sparing [Wells 2010], resulting in a tiny reduction of brain mass and a massive reduction in body mass. Second, during times of plenty, the body accumulates fat, whereas the brain does not. Seasonal variation in body mass may therefore greatly exceed the one in brain mass, except for a small number of species with seasonal variation in brain mass [Dechmann et al. 2017]. Third, among females body mass varies across the reproductive cycle, with higher values during pregnancy and lower values toward the end of lactation. Finally, captivity, too, can increase differences in body weight between wild and captive specimens, especially in slow-growing species [e.g. Isler et al. 2008; Leigh 1994]. Due to these various processes, variance in body mass at a given size can be up to four times as high as in brain mass [Pagel & Harvey 1989), and the slope will inevitably be underestimated, potentially by up to a factor two. To reduce this erroneous reduction in the slope estimate, we should include a greater range of intraspecific body sizes in the analysis, reduce the error in the estimation of each point by taking averages where possible, and use only wild specimens.

One seemingly obvious way to achieve this is to use the means of higher units (species, genera, families) as data points. This does indeed produce steeper slopes [Martin & Harvey 1985]. However, while this so-called taxon-level effect is partly due to the reduction of noise due to varying body mass, it also reflects the combined Cope-Marsh effects noted above, which will also produce higher slopes for families and orders [Jerison 1973; Rowe et al. 2011]. Therefore, because we cannot disentangle this taxon-level effect and the estimation error effect, slopes obtained among related species or genera need not reflect cognitive equivalence. However, this slope may still be useful, as it provides a convenient upper limit to the actual slope of cognitive equivalence. The analysis of congeneric slopes by Isler et al. [2008] using independent contrasts yielded a mean slope of 0.41 for primates, suggesting the true slope lies somewhere between 0.15 and 0.40.

The best option thus remains to obtain intraspecific slopes that are affected as little as possible by the error problem. We can achieve this by looking for species with large variation in adult body size. Using data on dog breeds, which show 30-fold variation in body size, Bronson [1979] obtained a slope of 0.27 based on averages per breed. This would seem to provide an excellent estimate of the slope of cognitive equivalence, but since dog breeds were produced by artificial selection, we cannot be sure that cognitive equivalence was maintained across the whole size range [cf. Martin & Harvey 1985].

Here, we consider a variation on this theme. We focus on primates with clear sexual dimorphism in body size, so as to minimize error due to the small range in adult body sizes. Second, we take the means of males and females so as to reduce the error in individual data points. And third, we only consider data from wild specimens. We then estimate the slope by fitting a line through the female and male average (which we will refer to as the 2-point slope). We predict that this 2-point slope will be steeper than the slope through all available data points, and closer to the true, unbiased value.

This 2-point slope will be biased when the sexes differ in body composition, in which case body mass is not a good estimate of actual (lean) body size [Schoenemann 2004]. In humans (not in our sample), men’s bodies are far leaner than women’s bodies, even among foragers [Wells 2010]. Obviously, the 2-point slope would then be overestimated. However, such major sex differences in adiposity are not found among non-human primates, because both arboreality and the high mobility under natural conditions will strongly limit adiposity and thus reduce any such differences [Heldstab et al. 2016; Sterck et al. 2019]. By including only animals taken from the wild, we largely eliminate this possible confounding effect.

The 2-point slope will also be biased when males and females experience differential selection pressures on overall cognitive abilities, in particular due to sexual selection, which by definition may affect the sexes differently. We do not expect this to be important because we know of no studies of primate cognition that needed to control for sex [e.g. Amici et al. 2012; Hopkins et al. 2014; Damerius et al. 2017; see also Arden & Adams 2016, for dogs]. Likewise, for humans most experts agree that there is no gender difference in intelligence, although some argue for a small difference [Irwing & Lynn 2005], which may also reflect differential socialization. Moreover, for primates, Lindenfors et al. [2007] found no evidence that sexual selection affected relative neocortex size, the largest part of primate brains. Thus, this assumption seems warranted. However, for birds, Garamszegi et al. [2005] found a weak positive effect of extra-pair copulation on female brain size. Likewise, Kotrschal et al. [2012] found that males in an Icelandic population of three-spined sticklebacks had far larger brains than females, although they could only speculate about the selective agent (perhaps male-only parental care). This effect is less likely in our sample because we focused on highly dimorphic species, thus excluding major variation in the mating system. Nonetheless, to control for the possibility, we will compare species with single-male versus multi-male mating. A final possible effect of sexual selection is that there may be an upper limit to the males’ ability to maintain cognitive equivalence as body mass dimorphism increases, because females are thought to be at the ecologically optimal size for a given niche. We therefore also examine the importance of dimorphism as a factor in the slope.

Given these various assumptions and our inability to fully test all of them, it is essential to seek external validation. Two opportunities exist. First, we can assess the ecological validity, by asking whether the residual brain size values based on the new slope actually predict cognitive abilities across species and do so better than alternative neuroanatomical measures that have been proposed in the past. This test faces a major hurdle, in that we do not know whether the relationship between the correct neuroanatomical measures and cognitive performance is linear, more than linear or less than linear. This problem is exacerbated by the fact that the performance measures are often normalized or even ordinal. However, since the relationship is necessarily monotonic, the correct measure should always preserve the rank order in cognitive performance. Although the different neuroanatomical measures we compare are likely to show high rank correlations among each other, we can nonetheless test their predictive value by assessing the value of the rank correlations with cognitive performance. We found two published data sets comparing species’ cognitive performance for this analysis. A second way to validate the slope value is to compare it with the strength of genetic correlations between brain size and body size [Lande 1979]. We will do this in the discussion, once we have acquired our estimate.

In this study, we therefore first determined the slopes of cognitive equivalence for sexually dimorphic primate species. We used two samples (one using conservative and another using more relaxed sample size criteria), and assessed the possible effects of sample sizes, mating system, and sexual dimorphism. Next, we validated the slope we obtained by comparing it with the genetic correlation between brain and body size and by its rank correlation with published measures of species’ cognitive abilities. Finally, we made a first, preliminary assessment of the consequences of adopting this new measure of a species’ cognitive abilities, using extinct hominins.

## Materialsand Methods

We compiled data on cranial capacity and body mass of primates from the studies by Heldstab et al. [2018, 2019]. We selected species with fairly large sample sizes (N ≥ 5 for each sex) to minimize error in estimating the mean body and brain size of males and females. We only included wild-caught animals to avoid captivity effects on body mass (usually fattening), and fully adult individuals, as evinced by the eruption of the third molar. Finally, we set a minimal mass dimorphism at 1.20, because preliminary analysis revealed wildly fluctuating estimated 2-point slopes at values close to monomorphism, as expected. Overall, we had 18 primate species that met these criteria. The brain-body size slope was then estimated for each species, as the slope of the line connecting the male and female average (the 2-point slope), as well as the slope through all points. A second primate data set was produced by including all species with at least 10 adult individuals and at least 2 of each sex and mass dimorphism of at least 1.20. This data set contained 27 species. We expected more variance in the estimate of the slope. We refer to these two data sets, as the conservative and relaxed primate data set, respectively.

We examined the bivariate effects of various variables discussed above on the estimated slope: mating system (single-male versus multi-male) and female brain size, as well as sexual dimorphism (to assess whether it has a positive effect on estimated slope for which we should control), sample size (likewise). We also used a model selection approach to identify the best-fitting model.

For the validation part, we used published data sets to capture cognitive performance: global cognitive performance [Deaner et al. 2007], and general intelligence factor *g*_*S*1_ from Reader et al. [2011].

The independent measures we used were:

1. body size (P, estimated as body mass in g);
2. brain size (E, estimated as endocranial volume, in cc, roughly corresponding to mass in g: Isler et al. 2008);
3. Jerison’s [1973] encephalization quotient (E/[0.12 x P^0.667^]),
4. An estimate of the cognitive brain based on cognitive equivalence (E- [0.065xP^0.27^]), where the exponent 0.27 is the midpoint value of the empirically obtained intraspecific 2-point slopes in the first set of analyses, and the intercept based on one of the smallest-brained mammals, *Sorex minutus*, which has a 5 gram body mass and a 0.1 gram brain weight [Bauchot & Stephan 1966]. This measure ensures that virtually all mammals have a positive cognitive brain size.

Note that measure #4 is an absolute measure of the amount of brain tissue available for cognitive tasks, whereas measure #3, the EQ, is a ratio. Thus, two species that differ greatly in body size can have the same EQ, yet the larger of the two will have far higher values of measure #4. Note, too, that measure #4 may be negative for ectothermic vertebrates [cf. Jerison 1973], and thus may only be intuitive for birds and mammals.

As noted above, we used rank correlations to assess the fit of these measures.

## Results

### Estimating the value of the slope of cognitive equivalence

As expected, the 2-point slopes based on mean values per sex are better than those based on all individual points. Using the conservative primate dataset, we found that the slopes using all the data points (mean= 0.209; SEM = 0.020) were on average 30% less steep than the 2-point slopes (mean= 0.268; SEM= 0.020). Likewise, the more conservative dataset, with at least 5 adult individuals of each sex, gave more reliable slope estimates (lower SEM) than the more relaxed data set, which also varied less with sexual dimorphism (Table 1). Thus, the best estimate of the slope is 0.27.

**Table 1.**
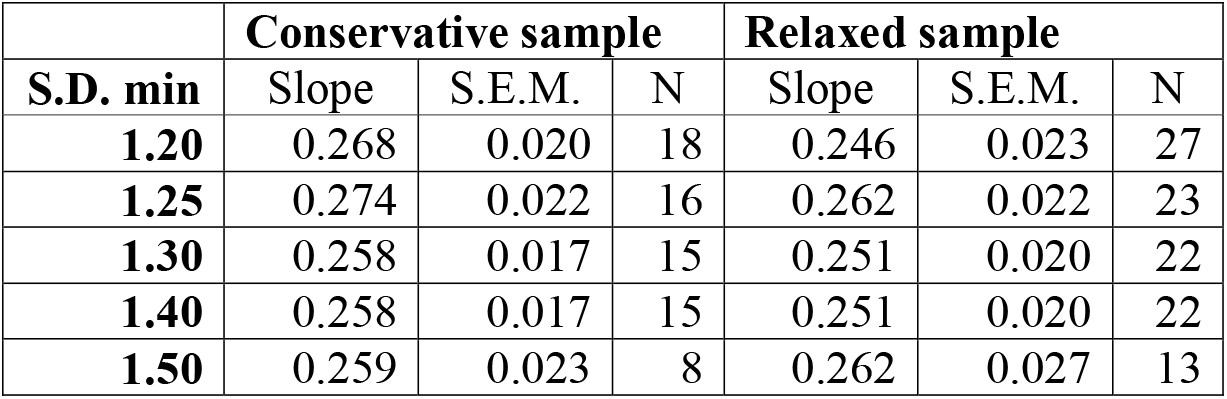
Effect of changing the minimal cutoff point of sexual dimorphism (S.D. min) for inclusion into the analysis of the 2-point slopes for the conservative and the relaxed primate sample (S.E.M. = standard error of the mean; N= number of species)

The analyses of the possible effect of confounding variables were performed on the conservative primate dataset only, again using only species with sexual dimorphism of ≥1.20, where the standard errors of the estimated slope values had stabilized. Our limited sample size forced us to do bivariate tests. In these tests, we did not need to correct for phylogenetic non-independence because the Pagel’s *λ* values of the slopes [Pagel 1992] were <0.001. We found no effect of sample size (r= −0.041, N= 18, P= 0.87) on the value of the 2-point slope. Likewise, there was no effect of sexual dimorphism (r= −0.093, p= 0.71) or of mating system (single-male versus multi-male; t_(16)_= −1.39, P= 0.18). Moreover, none of the various possible multivariate models showed anywhere near a significant result and multiple models had close similarity in overall fit (not shown).

### Validating the value of the slope

Table 2 provides the values of the Spearman rank correlations for the validation studies for each of the five measures used. Overall, as expected because of the need to use ranks only, the results are very close, and the confidence limits for the various measures generally overlap. Nonetheless, in both cognitive performance measures (the Deaner et al. [2007] and Reader et al. [2011] data), the EQ gives the lowest correlations, whereas the cognitive brain estimate, absolute brain size and even body size do about equally well.

**Table 2.** Validation analyses using multiple data sets of cognitive performance, using Spearman rank correlations.

| Predictor variable | $r_s$ | p-value | 95% CI | 99% CI |
| --- | --- | --- | --- | --- |
| *Deaner et al. (2007) N=24* |  |  |  |  |
| ln (brain size) | 0.811 | $p < 0.001$ | (0.507, 0.767) | (0.452, 0.795) |
| ln (body size) | 0.845 | $p < 0.001$ | (0.452, 0.736) | (0.393, 0.767) |
| cognitive brain | 0.809 | $p < 0.001$ | (0.602, 0.913) | (0.509, 0.933) |
| Jerison brain | 0.542 | $p = 0.007$ | (0.178, 0.775) | (0.045, 0.823) |
| *Reader et al. (2011) N=28* |  |  |  |  |
| ln (brain size) | 0.512 | $p = 0.005$ | (0.172, 0.742) | (0.050, 0.793) |
| ln (body size) | 0.515 | $p = 0.005$ | (0.176, 0.744) | (0.055, 0.794) |
| cognitive brain | 0.517 | $p = 0.005$ | (0.179, 0.746) | (0.057, 0.795) |
| Jerison brain | 0.408 | $p = 0.031$ | (0.042, 0.678) | (-0.081, 0.739) |

## Discussion

### Value of the slope

We examined the proposition that there is a slope of cognitive equivalence, which predicts the change in brain size, and thus cognitive abilities, corresponding to a change in body size in which all the details of bauplan and sensory-motor abilities are kept constant. Existing estimates of this slope suffer from the problem of measurement error, because mammals generally have only a narrow range of body sizes, whereas their estimates (body mass) can vary widely. We therefore used mean values for males and females in sexually dimorphic species, which have a larger range of body sizes, to estimate the 2-point slope. In general, this approach yielded steeper slopes than regressions through all available data points, confirming the expected reduction in the effect of error on slope estimates, and explaining why previous studies often found shallower slopes.

We found no effect of possible confounding variables, such as mating system or dimorphism beyond 1.20. The slope of 0.27 obtained here is the same as the one found for dogs, where breeds vary greatly in size [Bronson 1979], but, as expected, higher than the value of 0.23 [Hemmer 1971] or 0.25 [Röhrs 1986] found in previous intraspecific mammal studies with large samples that used all individual data rather than estimating 2-point slopes.

### Validation of the slope estimate

Based on selection experiments on body size in inbred mice lines, Lande [1979,1985] predicted a 0.36 slope of log (brain size) on log (body size), which would also hold when body size changes by drift, and thus also when populations contain selectively neutral genetic variation in body size. Slopes observed after selection experiments on body size in rodents yielded values between 0.2 and 0.4 [Riska & Atchley 1985], but since these should also be affected by the error problem, their true value may also be ≥ 0.25. However, Lande [1985] suggested that the genetic correlation between brain and body size is lower in natural species. Indeed, Grabowski [2016] estimated it for samples of five primate species taken from the wild, and found an average value of 0.254, which should be somewhat lower than the true slope, given the error in body mass used to estimate genetic correlation between brain and body. Thus, our estimate of 0.27 for the slope of cognitive equivalence is quite consistent with these estimates of genetic correlation.

We therefore used a line of P^0.27^ (where P is body size) as the estimate of the somatic brain. For estimates of a species’ cognitive brain we need to anchor the line (Fig. 1). To achieve positive values for all species, we took the presumably smallest-brained mammal, *Sorex minutus*, and forced the curve through its values (average body mass 5 g, average brain mass 0.1 g [Bauchot & Stephan 1966]). This leads to the minimum estimated size of the somatic brain as E_s_= 0.065 x P^0.27^, and thus the following estimate of the cognitive brain size: E_c_= E - E_s_, where E is the species’ actual brain size. Figure 1 illustrates the procedure.

**Figure 1.**
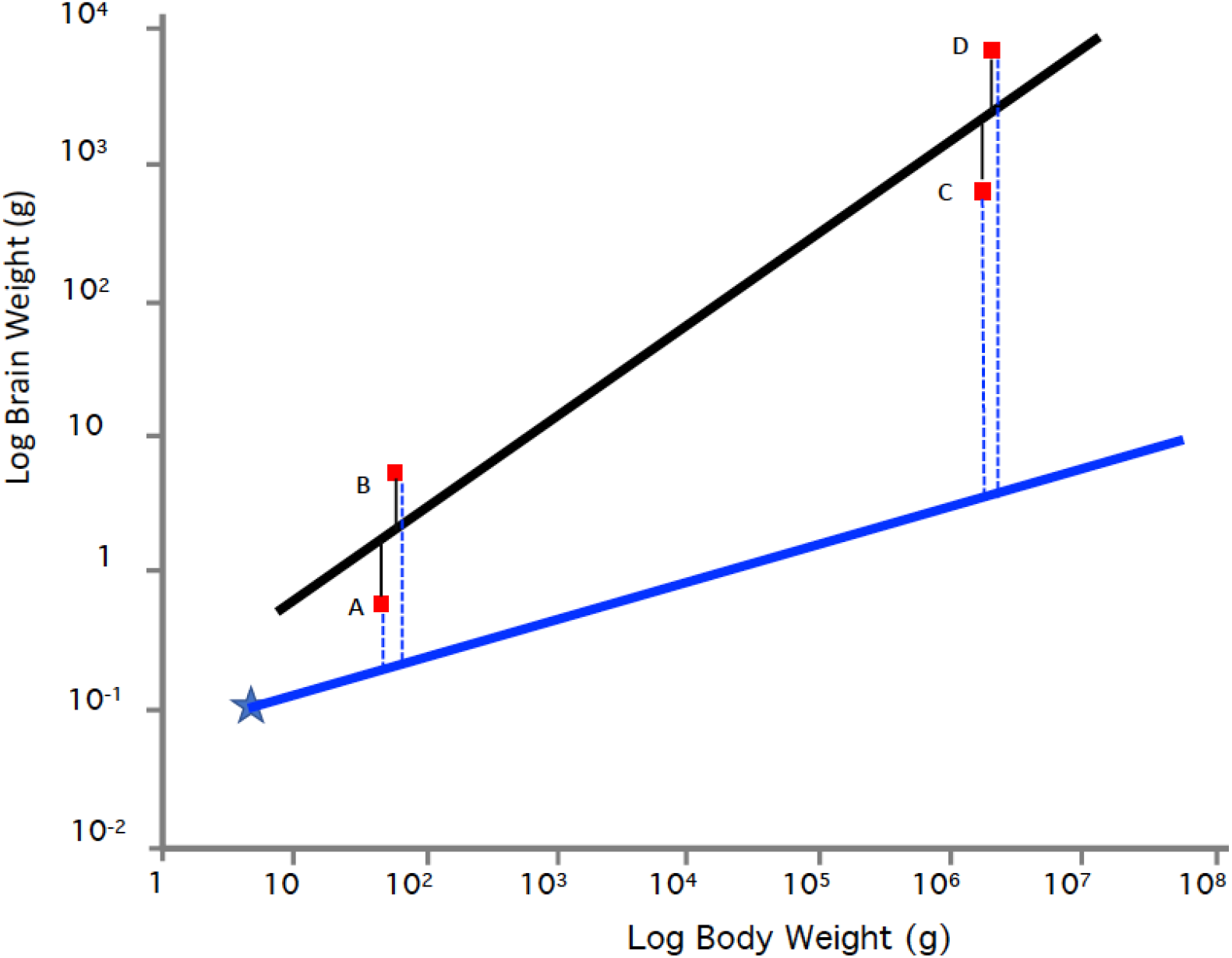
Illustrating the difference between EQs and estimates of cognitive brain size. The steep black line gives Jerison’s curve for the average mammalian brain and the shallow blue line the curve for the estimate of the somatic brain, which was anchored using the smallest-brained mammal, *Sorex minutus* (blue star). Four hypothetical species, red squares A-D, are included to illustrate the different measures. The black vertical lines are the residuals from Jerison’s line; EQ is the ratio of the value for a given point and the value of the corresponding point on the black line. The blue dashed lines are the estimates of the cognitive brain size.

The most important result of the cognitive validation analyses was that the alternative measures, including body size, all perform better than EQ. The EQ is the ratio of observed to expected brain size, and in Fig. 1, the antilog of the residual from Jerison’s line. Thus, a very small mammal could have the same ratio as a very large mammal (as do species B and D in Fig. 1), even if they differ dramatically in the absolute size of the estimated cognitive brain (the antilog of the blue dashed vertical lines). Thus, the estimated order of overall cognitive performance would increase as follows under the EQ: A=C < B=D. Under the cognitive brain estimate deployed here it would be: A < B < C <D, the same order as in absolute brain size, or(in this example) absolute body size of the species.

The close similarity in how well the various non-EQ measures predicted actual estimates of overall cognitive abilities is not surprising. Absolute brain size is often a reasonable predictor of specific cognitive abilities in studies that did not use the measure based on cognitive equivalence [Benson-Amran et al. 2015; MacLean et al. 2014; Horschler et al. 2019]. The two measures are closely correlated, and also of course show strong correlations with body size. Indeed, also the neocortex ratio remains highly correlated with body size [e.g. Stout 2018]. Although we did not include the neocortex ratio in this study, in the dataset used by Dunbar [1992], which uses averages for genera and sexes, the Spearman rank correlation between the two is 0.871 (N= 38, P<0.001) (non-linear relationship). Thus, the discrepancies among the various alternative approaches (absolute brain size, cognitive equivalence, neocortex ratio) are quite modest, and in practice we will often not have the resolution needed to decide which of these is the best.

Nonetheless, although the low value of the slope of cognitive equivalence indicates that only a modest correction for body size is needed, some correction for the somatic functions of brains is conceptually necessary (despite proposals to abandon it altogether). First, the cognitive abilities of extremely large animals would almost certainly be overestimated. Second, sex differences in cognitive abilities would otherwise be inferred that may well not exist. For instance, despite a clear gender difference in brain size [e.g. in Martin [1986], men’s mean is 1498 cc and women’s mean 1326 cc), there is only a tiny (if any) gender difference in intelligence in humans [Irwing & Lynn 2005]. The estimate based on cognitive equivalence is therefore to be preferred.

It appears that this approach reduces the effect of body size on brain size much more than has traditionally been considered necessary. It is not clear why this is. First, this conclusion may be peculiar to primates, with their large brains for mammals, and thus large neocortices [Finlay et al. 2001]. Where cognitive functions take up a large proportion of the overall brain size, the relative importance of the somatic functions must become concomitantly smaller. If so, we expect the slope to be lower for other mammalian orders. Second, the size of the somatic brain may be small overall because neurons in many brain regions can be involved in multiple networks simultaneously, thus blurring the distinction between somatic and cognitive functions.

The analysis presented here is of course preliminary. It is based on primates only, and on a limited number of species at that, and could not establish whether species differences in slopes were true or merely reflected error. Future work should include all mammals with non-trivial dimorphism in body size, while ensuring the use of fully adult, wild specimens, with values not affected by advanced pregnancy or confounded by seasonal variation in adiposity. Along with work on other vertebrates, this work may find that slopes vary among lineages, and generate and test hypotheses on its causes.

Nevertheless, this study confirms that the effect of body size on the size of the somatic brain (the 2-point slope) varies considerably between mammals (and presumably birds [Tsuboi et al. 2018]) on the one hand, and ectothermic vertebrates (fishes, amphibians, reptiles) on the other hand. The latter vary between 0.4 and 0.5 within species (Tsuboi et al 2018). For fishes, Triki et al. [2021, preprint] obtained a mean intraspecific slope of 0.49 that is independent of body size variation. Furthermore, Triki et al. [2021, preprint] studied one fish species, the cleaner fish *Labroides dimidiatus* in more detail. Individual performance in various cognitive tasks was not correlated with body size, so that the intraspecific brain-body slope of 0.53 of this species indeed represents the slope of cognitive equivalence. The causes of this clear difference between intraspecific slopes in endotherms versus ectotherm vertebrates remain unexplained.

It might be objected that comparing distantly related taxa on their cognitive performance based on brain measures only is not advised given the known differences in neuron densities in the brains of different clades [Herculano-Houzel 2017] and the effect on brain size of highly divergent bauplans. We fully endorse this view. Comparing cognitive brain estimates among distantly related lineages may not be very revealing, and we should be very careful. Nonetheless, this restriction does not argue in favor of reviving the EQ approach.

### Applications: An example

Although a thorough application of the new method is beyond the scope of this paper, we can get a first impression by comparing the cognitive brain estimate with Jerison’s [1973] EQ for extinct hominins thought to be in the main lineage contributing representatives that formed our own species (thus excluding the robust australopithecines and both *Homo floresiensis* and *H. naledi*). Because the new measure produces cognitive brain size estimates rather close to overall brain size, the two techniques yield a rather different picture (Figure 2). Jerison’s EQ suggests a long period of a continuing, gradual increase in cognitive abilities from 4 Ma until roughly 300 ka, followed by a sudden uptick. The alternative EQ measure proposed by Grabowski et al. [2016], which is based on a somewhat shallower slope, gives a similar picture, but without the sudden uptick in the last 300 ka (see their Fig. 2). The cognitive equivalence measure, in contrast, suggests cognitive stasis between 4 Ma and 2 Ma, after which cognitive abilities steadily increased.

**Figure 2.**
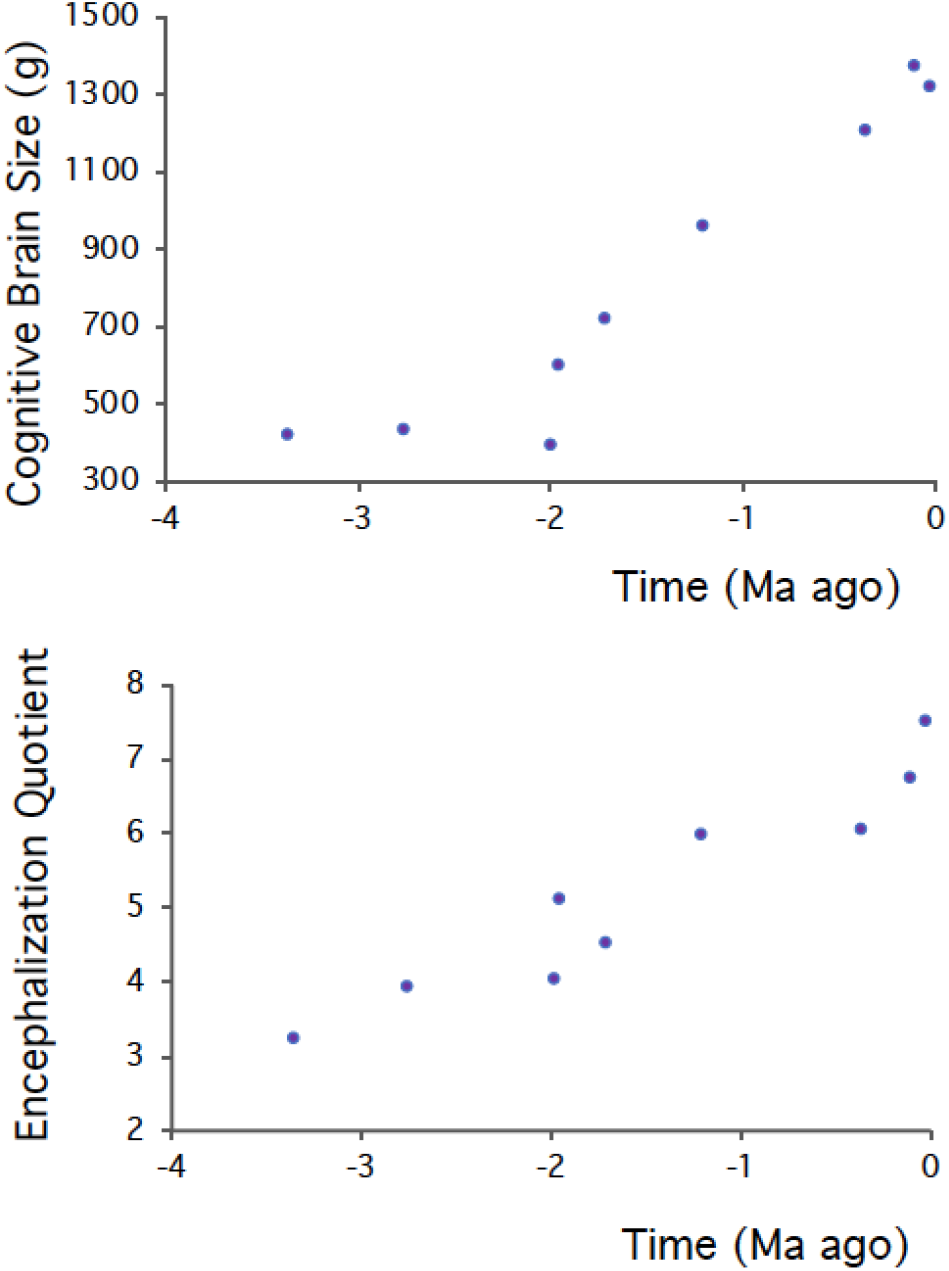
The changes in cognitive brain size (a) and EQ (b) during hominin evolution. The dates are the median observed age (Ma before present). Data based on Grabowski [2016].

The paucity of material and the problems inherent in estimating body size from incomplete remains advise caution. Still, it can be argued that the general pattern suggested by the cognitive equivalence measure is closer to current understanding. Thus, the material culture of hominins did not exceed that of the extant great apes [Wynn et al. 2011], the diversity and complexity of hominin material culture began in earnest around 2 Ma [e.g. Stout 2011], and there is no great increase in relative brain size within the human species [Grabowski et al. 2016; Schoenemann 2013].

### Implications

Comparative tests of adaptive hypotheses to explain brain size evolution in primates sometimes reach incompatible conclusions [Wartel et al. 2019], as do those on other mammals or birds [Healy & Rowe 2007]. Among the many reasons responsible for this impasse, one potentially prominent aspect is generally overlooked [Rogell et al. 2019]. These tests invariably include body mass as a control variable, and then try to explain the remaining variation in brain size, in effect letting the pattern of covariation among the variables decide the allometric slope. In practice, the high correlation between body size and the other potential explanatory variables may affect the outcome and interpretation of the results [Stout 2018].

Thus, when we have a validated estimate of the size of the cognitive brain we can use this residual measure as the response variable, and so should get a fairer evaluation of the hypotheses trying to explain brain size evolution. In the end, this may be the most valuable contribution of the kind of analysis undertaken in this paper.

## Acknowledgments

We thank Tsuboi Masahito for comments on an earlier version of the manuscript. CvS thanks the Eugene Dubois Foundation for supporting his stay in Maastricht, which inspired him to take up the quest again.

## Ethics note

Data were compiled from published studies.

## Authors’ contributions

CvS provided the conceptual basis, and RB and ZT added major discussion; SH provided data; SH and CvS did the analyses; all authors wrote the final version.

